# RCVT: a diagnostic to quantify compositional biases among taxa for large phylogenomic studies

**DOI:** 10.1101/2024.11.28.625917

**Authors:** Jacob L Steenwyk, Thomas J. Buida

## Abstract

Phylogenomics aims to reconstruct the history of genes and genomes. However, noise or error during inference can stem from diverse sources, such as compositional biases. Here, we introduce RCVT (**<u>R</u>**elative **<u>C</u>**omposition **<u>V</u>**ariability among **<u>T</u>**axa), a metric to quantify compositional biases among taxa. We demonstrate the utility of RCVT using example data and quantify compositional biases in 16 empirical phylogenomic datasets, revealing variation in bias among taxa within phylogenomic data matrices. Systematic removal of taxa with high RCVT scores substantially reduces compositional heterogeneity compared to randomly pruning taxa among large phylogenomic data matrices. RCVT may help researchers diagnose and potentially ameliorate phylogenomic noise associated with compositional biases.

## Introduction

Phylogenetics aims to reconstruct the evolutionary history of genomes, genes, and other biological entities (Kapli et al. 2020; Steenwyk et al. 2023). However, sources of noise and error can cause inaccurate phylogenetic inferences (Betancur-R. et al. 2019; Steenwyk et al. 2023). Accordingly, numerous metrics have been devised to diagnose potential sources of error at the level of taxa, genes, and sites. For example, there are numerous metrics to diagnose potential errors among genes, such as alignment length, the number of parsimony informative sites, saturation, and clock-like evolution (Philippe et al. 2011; Shen et al. 2016; Liu et al. 2017; Steenwyk et al. 2020). In contrast, there are fewer metrics to evaluate compositional bias among taxa, though they have demonstrated that compositional biases may be widespread in phylogenomic studies (Jermiin et al. 2004; Naser-Khdour et al. 2019; Smith et al. 2023).

Here, we introduce a metric to quantify compositional bias among taxa termed RCVT (**<u>R</u>**elative **<u>C</u>**omposition **<u>V</u>**ariability among **<u>T</u>**axa). To demonstrate the utility of this metric, we calculate RCVT in a fictitious four-taxon case. Thereafter, we calculate RCVT in 16 phylogenomic data matrices from diverse lineages and uncover variation in compositional biases among taxa therein. In 15 out of the 16 datasets, taxa with outlier RCVT values were identified, which suggests that sequences with outlier compositional biases are prevalent and may warrant further scrutiny in phylogenomic studies, corroborating previous findings (Naser-Khdour et al. 2019). We also demonstrate that RCVT-informed taxon pruning strategies reduce compositional heterogeneity in large phylogenomic datasets, while smaller datasets have a less consistent reduction in compositional heterogeneity.

To facilitate researchers calculating RCVT, we incorporate a new function to calculate RCVT in PhyKIT, a command-line toolkit for processing and analyzing phylogenomic data (i.e., multiple sequence alignments and phylogenetic trees) (https://jlsteenwyk.com/PhyKIT) (Steenwyk et al. 2021, 2024).

## Methods

RCVT measures how much a particular taxon differs from the average composition of all taxa in a multiple-sequence alignment. RCVT is adapted from the metric *relative composition variability*, which measures compositional bias at the gene level (Phillips and Penny 2003). More specifically, RCV is the average compositional variability among taxa, whereas RCVT is the per taxon compositional variability values used to calculate RCV. The lower limit of RCVT values is 0 representing low compositional bias; high values indicate greater compositional biases. The following formula is used to calculate RCVT:

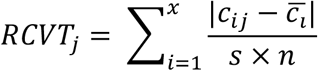

Wherein abbreviations are as follows:

- *RCVT*_*j*_: relative composition variability for taxon *j*, which quantifies its variability compared to others;
- *c*: the number of different character states per sequence type (e.g., A, T, C, and G for DNA sequences) in an alignment;
- 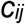: the number of the *i*th character state (e.g., how many times A appears) for the *j*th taxon;
- 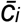: the average number of the *i*th *c* character state across *n* taxa, which represents the general occurrence of that character in a dataset;
- *s*: total number of sites; and
- *n*: the number of taxa.

Using this formula, higher values are associated with greater compositional biases, whereas lower values are associated with less bias. For example, in a toy four-taxon case, one taxon may have only one nucleotide character, but the other three taxa have an equal number of each of the four nucleotides (A, T, C, and G). In this case, the taxon with only one nucleotide character will have a much higher RCVT than the other three taxa (Figure 1a).

**Figure 1.**
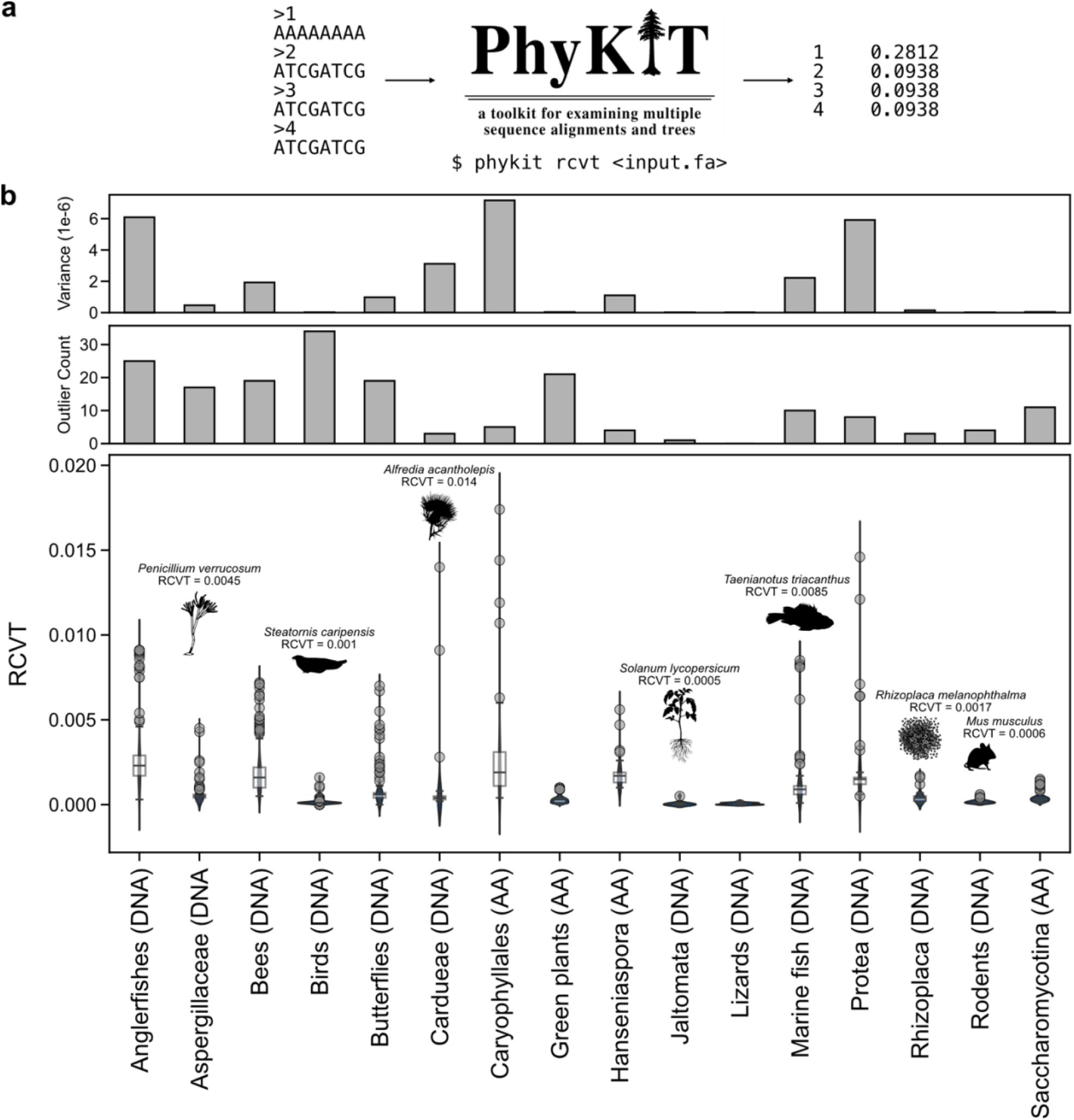
RCVT measures compositional bias and reveals that compositional outliers are common among phylogenomic datasets. (a) In a toy dataset wherein a taxon has only one character state in an alignment, whereas the other taxa have equal numbers of the four nucleotides, the taxon with one character state has a higher RCVT value. RCVT can easily be calculated using the *rcvt* function in the command-line tool, PhyKIT (Steenwyk et al. 2021). (b) The distribution of individual RCVT values reveals variation in compositional biases among 16 empirical datasets (Table 1). (b, top) Each dataset had different degrees of variance among RCVT values and (b, middle) different numbers of outlier taxa. (b, bottom) Boxplots with violin plots provide additional information about the median, 75^th^, and 25^th^ percentiles. For select datasets, we highlight outlier taxa and their RCVT values; for example, in the Aspergillaceae dataset, *Penicillium verrucosum* had the highest RCVT value of 0.0045. Note that the lower limit of RCVT is zero, and although it appears that some values fall below zero in the violin plot, they are an artifact of the kernel density estimation of the underlying distribution. The character type (DNA or amino acids [AA]) are specified after the dataset name.

A function to calculate RCVT has been incorporated into the Python-based tool, PhyKIT (Steenwyk et al. 2021), as of version 1.13.0 (https://jlsteenwyk.com/PhyKIT/usage/index.html#relative-composition-variability-taxon). To ensure the efficacy of the newly added function, we wrote five integration tests that cover 100% of the code for the RCVT function. These tests, alongside a continuous integration pipeline, which leverages architecture built elsewhere (Steenwyk et al. 2020, 2022b,a; Steenwyk and Rokas 2021), helps ensure PhyKIT correctly builds, packages, and installs across multiple versions of Python. These efforts were motivated by nearly 30% of all bioinformatic tools failing to install (Mangul et al. 2019), which threatens the reproducibility of bioinformatics research. More broadly, we aim to provide the research community with a long-term dependable tool for phylogenomics research.

**Table 1.**
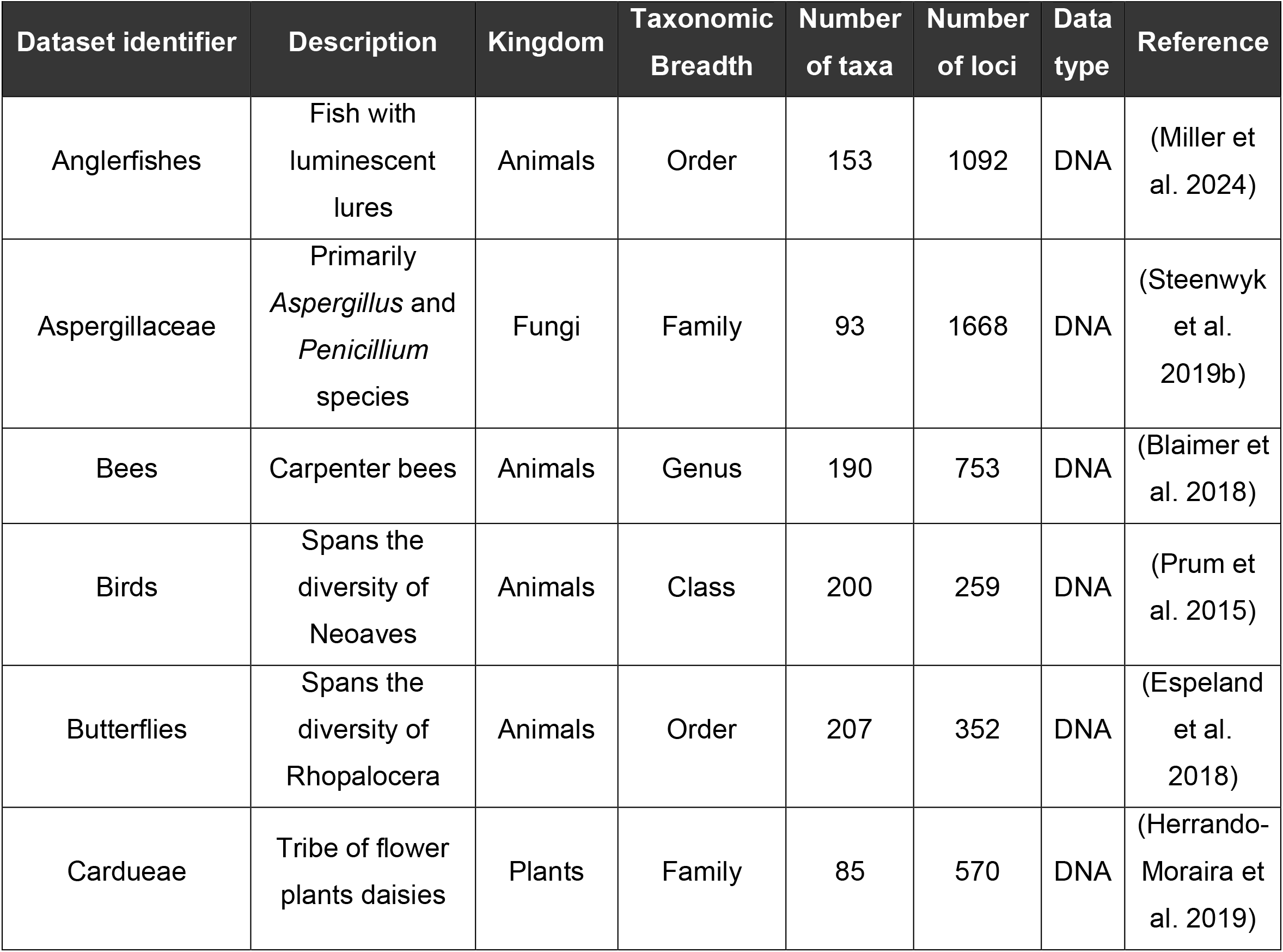
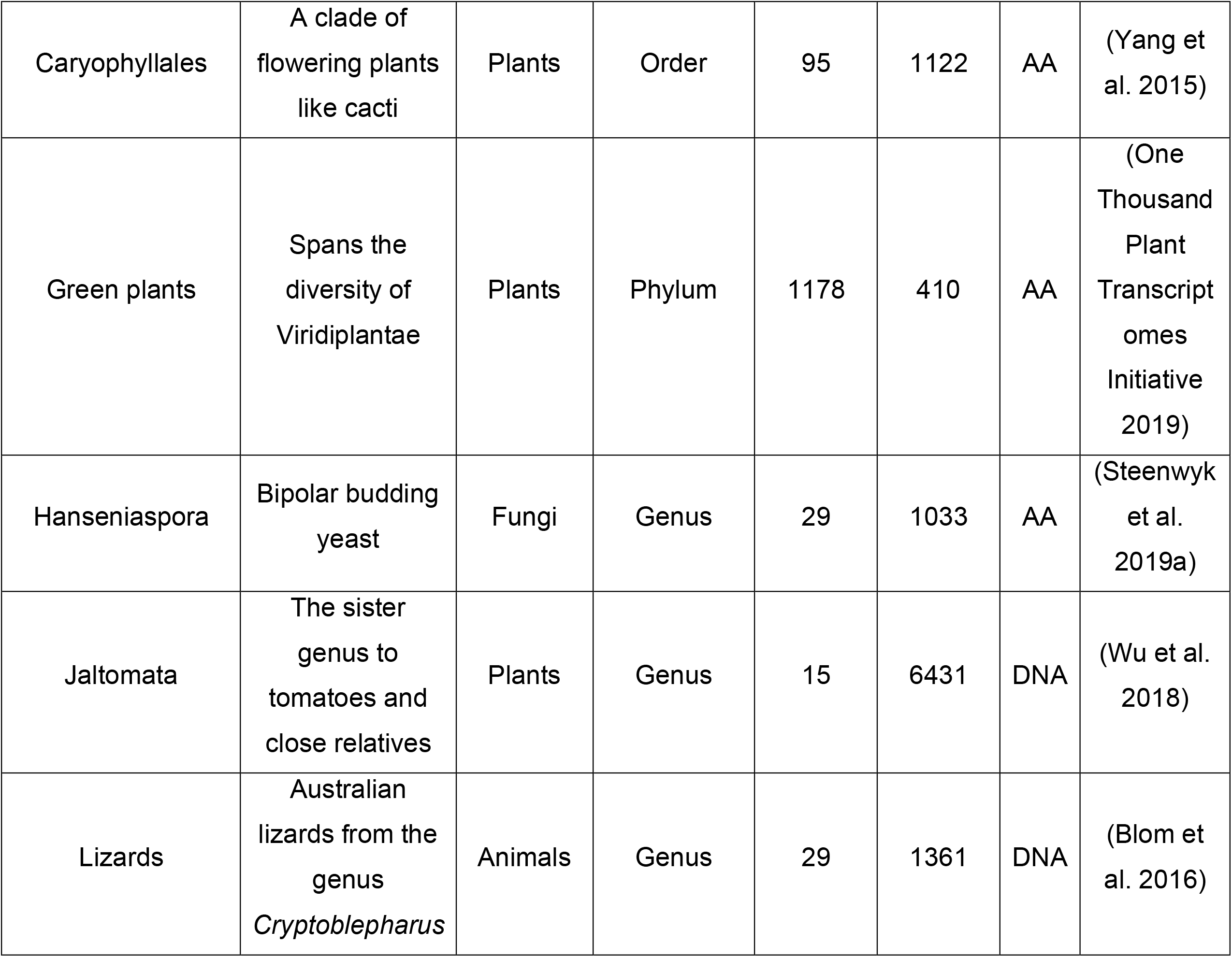
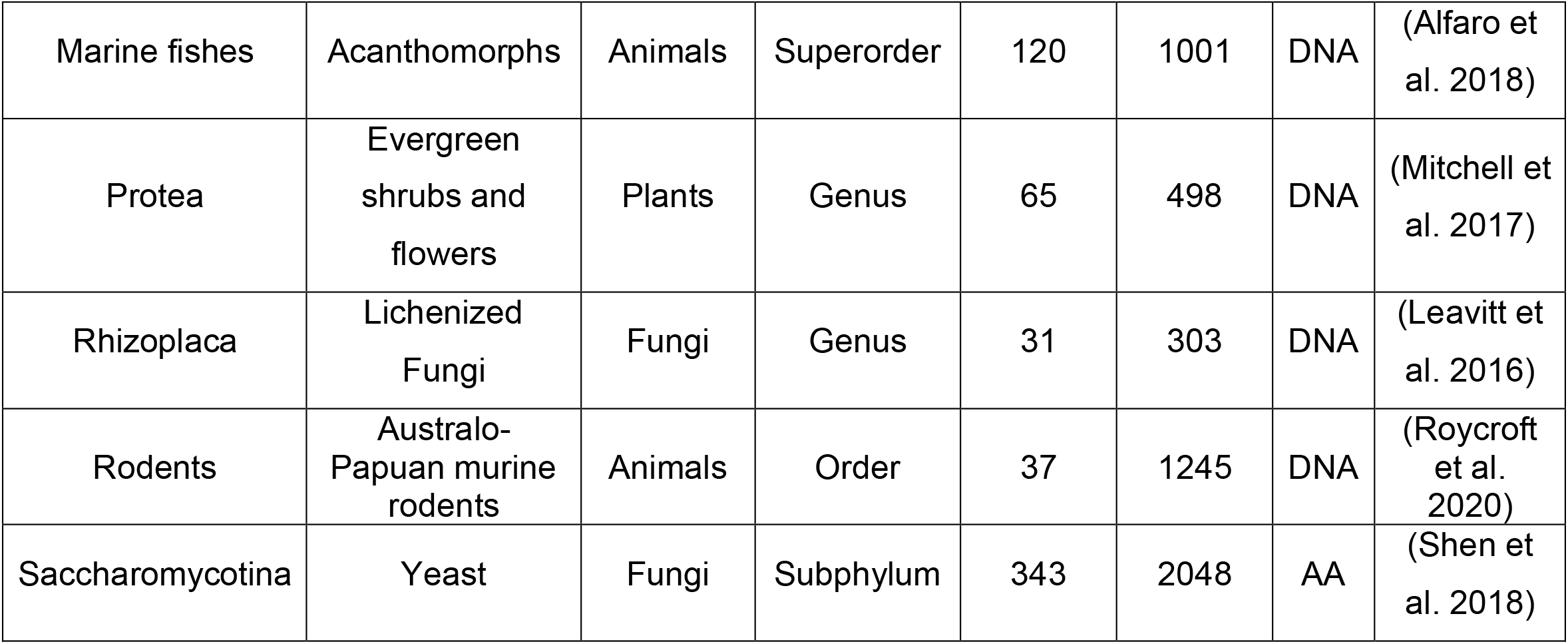
Empirical phylogenomic datasets examined.

We also calculated RCVT for 16 empirical phylogenomic datasets (Table 1). For each dataset, single-gene multiple sequence alignments were downloaded from a previous study that examined phylogenomic methodology (Liu et al. 2024) or from the primary study, as in the case of the Anglerfishes dataset (Miller et al. 2024). The resulting multiple sequence alignments were then concatenated into a supermatrix using the *create_concat* function in PhyKIT (Steenwyk et al. 2021). Next, RCVT was calculated for the taxa in each supermatrix using the *rcvt* function in PhyKIT. To detect outlier values, we identified RCVT values that had values greater than the 75^th^ percentile plus

1.5 times the interquartile range (the difference between the 75^th^ and 25^th^ percentiles).

We also evaluated the impact of removing taxa with high RCVT values on compositional heterogeneity. To do so, one to *N* taxa—wherein *N* represents the total number of taxa in a data matrix—with the highest RCVT values were removed with a step of one from each data matrix. Thereafter, χ^2^ statistics were calculated to measure compositional heterogeneity; higher values indicate higher compositional heterogeneity, and lower values indicate lower heterogeneity (or that all sequences are more similar). To generate a null distribution, *N* number of randomly selected taxa were removed and χ^2^ statistics were recalculated. This process was done 10 times. The mean and 95% confidence intervals were calculated from the 10 iterations.

## Results and Discussion

To determine how common compositional outliers are, we calculated RCVT in 16 phylogenomic datasets (Figure 1b and Table 1). These datasets are composed of amino acid or nucleotide sequences, span a diversity of taxonomic ranks, lineages, number of taxa, and loci. Quantifying the number of taxa with high outlier RCVT values revealed variation in the number of outlier taxa per dataset (Figure 1b, middle panel). For example, the dataset of birds had the highest number of outliers. Similarly, the anglerfish dataset also had many outliers. These findings corroborate previous findings that compositional heterogeneity is widespread in phylogenomic datasets (Jermiin et al. 2004; Naser-Khdour et al. 2019).

In contrast, other datasets had few taxa with outlier RCVT values. For example, the dataset of Rodents and Saccharomycotina yeast had few outliers. The dataset of Lizards had no outlier taxa. Examination of the taxa that were outliers revealed, at times, that outlier taxa were outgroup taxa. There was no obvious association with sequences type—nucleotides or amino acids—and the likelihood of there being outliers.

Next, we determined if an RCVT-informed taxon pruning strategy helps reduce compositional heterogeneity. To do so, we removed one to *N* taxa—wherein *N* is the total number of taxa in a data matrix—with the highest RCVT scores in each data matrix and examined compositional heterogeneity, which was measured using the χ^2^ statistic. A null distribution was generated by randomly pruning one to *N* taxa from a data matrix 10 times and taking the resulting mean and 95% confidence interval from the calculated χ^2^ statistics.

Examination of the resulting distributions revealed RCVT-informed taxon pruning reduced compositional heterogeneity in large data matrices (approximately 100 or more taxa; Figure 2). Among smaller datasets, RCVT-based pruning did not consistently reduce compositional heterogeneity; for example, in the Aspergillaceae dataset compositional heterogeneity was reduced for approximately the first 25 taxa, while in the Rodents datasets, RCVT-based pruning is not substantially different from the null distribution.

**Figure 2.**
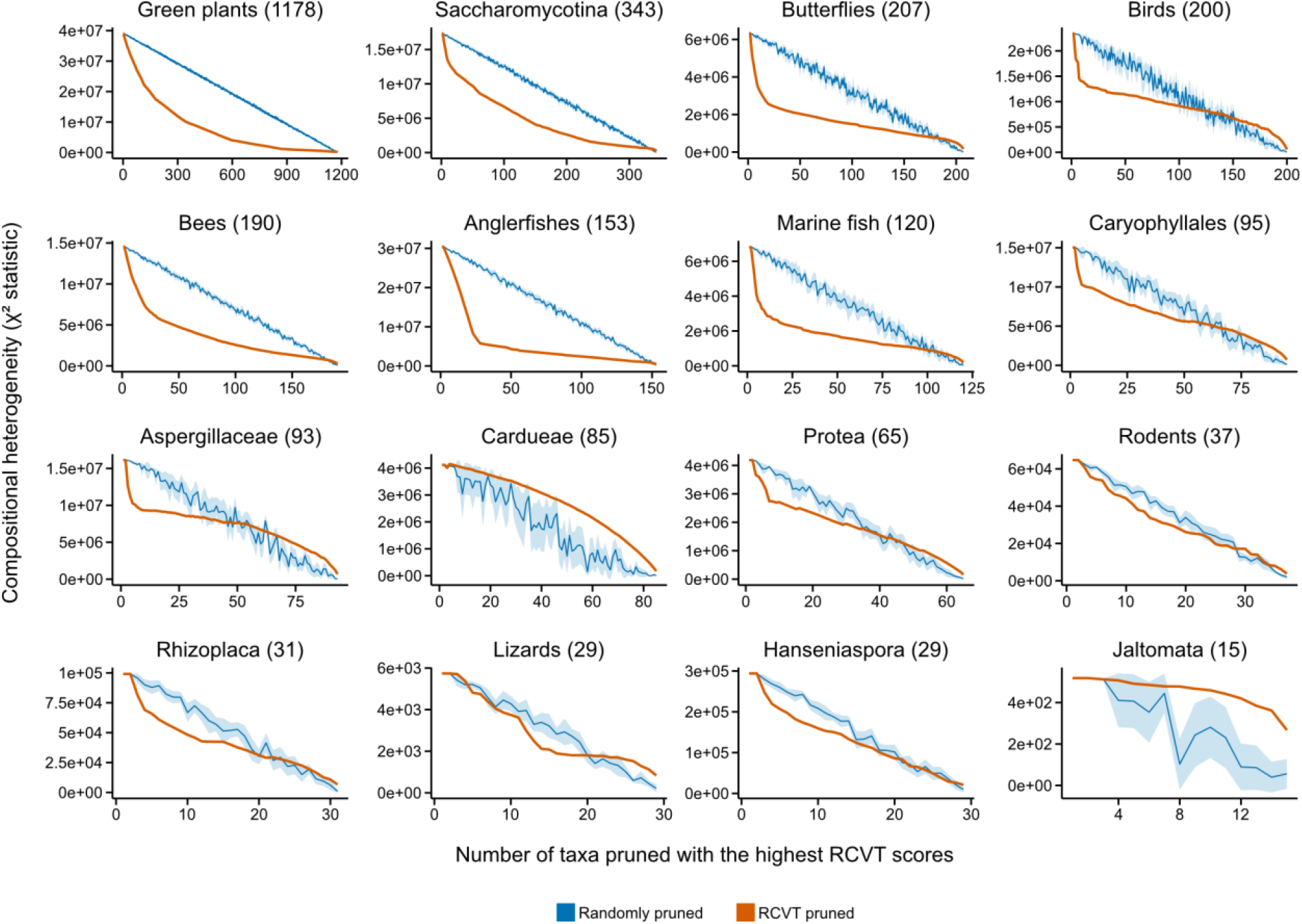
Removing taxa with high RCVT scores reduces compositional heterogeneity among large phylogenomic datasets. Each panel depicts compositional heterogeneity (quantified as the χ^2^ statistic; y-axis) wherein lower values indicate less heterogeneity with a different number of taxa pruned from the dataset (x- axis). To create a null distribution, random sets of taxa of *N* taxa (where *N* is one to the number of total taxa in a dataset) were pruned 10 times and the χ^2^ statistic was calculated for each iteration; the line represents the average χ^2^ statistic with a 95% confidence interval (blue). To evaluate if pruning using an RCVT-informed strategy lowers compositional heterogeneity better than randomly selected taxa, the top *N* number of taxa with the highest RCVT scores were removed from a dataset and the χ^2^ statistic was recalculated (orange). Examination of the resulting distributions revealed that χ^2^ statistic scores were lower when pruning taxa based on RCVT scores, namely, among large phylogenomic datasets (approximately 100 or more taxa). Smaller datasets (approximately fewer than 100 taxa) did not benefit as much from the RCVT- based pruning strategy. The number of taxa in each dataset is next to the dataset name in parentheses. Panels are arranged from the largest (top left) to the smallest (lower right) dataset size based on the number of taxa.

These findings suggest that compositional biases may be prevalent in phylogenomic data matrices and that large data matrices of 100 or more taxa had the most apparent reduction in compositional biases when pruning taxa with high RCVT values. RCVT, alongside other measures of compositional bias (Naser-Khdour et al. 2019), may help explore—and quantify—compositional biases among taxa in large phylogenomic data matrices.

### Practical considerations

The intended use of RCVT is to identify individual taxa with putative compositional biases (Figure 1) that could negatively impact phylogenomic inference. After removing taxa with high RCVT values across each empirical dataset, examination of compositional heterogeneity revealed that large phylogenomic data matrices (approximately 100 or more taxa) had the most evident reduction in compositional heterogeneity (Figure 2). Thus, RCVT may be most useful for large phylogenomic data matrices.

RCVT may be integrated into the phylogenomic subsampling approach to explore stability of inferences (Edwards 2016). In phylogenomic subsampling, taxa (or genes) are subsampled according to diverse metrics. For example, gene-based strategies may subsample genes with the highest most parsimony informative sites because they are associated with higher accuracy and confidence during single-gene phylogenetic inference (Shen et al. 2016; Steenwyk et al. 2020). Thereafter, a phylogenomic tree is inferred using a subset of the total phylogenomic data matrix, and the resulting phylogenomic tree is compared to the tree inferred from the entire data matrix, identifying potentially unstable branches. Analogously, taxa with high RCVT scores (for example, the top 5% of taxa) may be pruned from a dataset, and the phylogenomic tree can be inferred and compared to the full taxon data matrix to identify branches susceptible to compositional heterogeneity.

Though our analyses suggest RCVT may help diagnose taxa that may contribute to phylogenomic error through compositional biases, RCVT is a novel metric and, therefore, warrants additional exploration and scrutiny. One potential avenue for exploration is to remove the *j*th taxon when calculating *c*_*ij*_ and 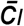. By excluding sequence *j* from the character-frequency calculations, it may be easier to determine whether that sequence is an outlier. Furthermore, RCVT does not consider site homology, unlike other tests such as the matched-pairs homogeneity tests, or evolutionary events (Bowker 1948; Stuart 1955; Ababneh et al. 2006; Naser-Khdour et al. 2019; Smith et al. 2023). Adapting RCVT to account for site-wise homology, evolutionary events, or other factors may improve the utility of the metric. Nonetheless, RCVT, alongside other tests, may be used as a complementary method to explore potential compositional biases.

Another potential difficulty in interpreting RCVT values is that the null distribution for RCVT values is complex to discern (Posada 2009). Here, we used a simple outlier detection method based on interquartile ranges; however, more statistically complex methods may be applied to identify outliers. Also, since RCVT is a relative metric, it is not advised to compare RCVT values between data matrices. Instead, RCVT values are best compared within a given data matrix.

## Data availability

RCVT has been incorporated into PhyKIT (Steenwyk et al. 2021) and comes complete with documentation (https://jlsteenwyk.com/PhyKIT/usage/index.html#relative-composition-variability-taxon). PhyKIT is freely available under the MIT license and can be downloaded from PyPi (https://pypi.org/project/phykit/) or GitHub (https://github.com/JLSteenwyk/PhyKIT).

## Acknowledgements

JLS thanks Kassiani Panagiotou for the helpful discussion while at a workshop together.

## References

Ababneh, F., L. S. Jermiin, C. Ma, and J. Robinson . 2006. Matched-pairs tests of homogeneity with applications to homologous nucleotide sequences. Bioinformatics 22:1225–1231.

Alfaro, M. E., B. C. Faircloth, R. C. Harrington, L. Sorenson, M. Friedman, C. E. Thacker, C. H. Oliveros, D. Černý, and T. J. Near. 2018. Explosive diversification of marine fishes at the Cretaceous–Palaeogene boundary. Nat Ecol Evol 2:688–696.

Betancur-R., R., D. Arcila, R. P. Vari, L. C. Hughes, C. Oliveira, M. H. Sabaj, and G. Ortí. 2019. Phylogenomic incongruence, hypothesis testing, and taxonomic sampling. Evolution 73:329–345. [Society for the Study of Evolution, Wiley].

Blaimer, B. B., J. R. Mawdsley, and S. G. Brady. 2018. Multiple origins of sexual dichromatism and aposematism within large carpenter bees. Evolution 72:1874–1889.

Blom, M. P. K., J. G. Bragg, S. Potter, and C. Moritz. 2016. Accounting for Uncertainty in Gene Tree Estimation: Summary-Coalescent Species Tree Inference in a Challenging Radiation of Australian Lizards. Syst Biol syw089.

Bowker, A. H. 1948. A Test for Symmetry in Contingency Tables. Journal of the American Statistical Association 43:572–574.

Edwards, S. V. 2016. Phylogenomic subsampling: a brief review. Zool Scr 45:63–74.

Espeland, M., J. Breinholt, K. R. Willmott, A. D. Warren, R. Vila, E. F. A. Toussaint, S. C. Maunsell, K. Aduse-Poku, G. Talavera, R. Eastwood, M. A. Jarzyna, R. Guralnick, D. J. Lohman, N. E. Pierce, and A. Y. Kawahara. 2018. A Comprehensive and Dated Phylogenomic Analysis of Butterflies. Current Biology 28:770-778.e5.

Herrando-Moraira, S., J. A. Calleja, M. Galbany-Casals, N. Garcia-Jacas, J.-Q. Liu, J. López-Alvarado, J. López-Pujol, J. R. Mandel, S. Massó, N. Montes-Moreno, C. Roquet, L. Sáez, A. Sennikov, A. Susanna, and R. Vilatersana. 2019. Nuclear and plastid DNA phylogeny of tribe Cardueae (Compositae) with Hyb-Seq data: A new subtribal classification and a temporal diversification framework. Molecular Phylogenetics and Evolution 137:313–332.

Jermiin, L. S., S. Y. W. Ho, F. Ababneh, J. Robinson, and A. W. D. Larkum. 2004. The Biasing Effect of Compositional Heterogeneity on Phylogenetic Estimates May be Underestimated. Systematic Biology 53:638–643.

Kapli, P., Z. Yang, and M. J. Telford. 2020. Phylogenetic tree building in the genomic age. Nat Rev Genet 21:428–444.

Leavitt, S. D., F. Grewe, T. Widhelm, L. Muggia, B. Wray, and H. T. Lumbsch. 2016. Resolving evolutionary relationships in lichen-forming fungi using diverse phylogenomic datasets and analytical approaches. Sci Rep 6:22262.

Liu, C., X. Zhou, Y. Li, C. T. Hittinger, R. Pan, J. Huang, X. Chen, A. Rokas, Y. Chen, and X.-X. Shen. 2024. The Influence of the Number of Tree Searches on Maximum Likelihood Inference in Phylogenomics. Systematic Biology 73:807–822.

Liu, L., J. Zhang, F. E. Rheindt, F. Lei, Y. Qu, Y. Wang, Y. Zhang, C. Sullivan, W. Nie, J. Wang, F. Yang, J. Chen, S. V. Edwards, J. Meng, and S. Wu. 2017. Genomic evidence reveals a radiation of placental mammals uninterrupted by the KPg boundary. Proc. Natl. Acad. Sci. U.S.A. 114.

Mangul, S., T. Mosqueiro, R. J. Abdill, D. Duong, K. Mitchell, V. Sarwal, B. Hill, J. Brito, R. J. Littman, B. Statz, A. K.-M. Lam, G. Dayama, L. Grieneisen, L. S. Martin, J. Flint, E. Eskin, and R. Blekhman. 2019. Challenges and recommendations to improve the installability and archival stability of omics computational tools. PLoS Biol 17:e3000333.

Miller, E. C., R. Faucher, P. B. Hart, M. Rincón-Sandoval, A. Santaquiteria, W. T. White, C. C. Baldwin, M. Miya, R. Betancur-R, L. Tornabene, K. Evans, and D. Arcila. 2024. Reduced evolutionary constraint accompanies ongoing radiation in deep-sea anglerfishes. Nat Ecol Evol, doi: 10.1038/s41559-024-02586-3.

Mitchell, N., P. O. Lewis, E. M. Lemmon, A. R. Lemmon, and K. E. Holsinger. 2017. Anchored phylogenomics improves the resolution of evolutionary relationships in the rapid radiation of Protea L. American J of Botany 104:102–115.

Naser-Khdour, S., B. Q. Minh, W. Zhang, E. A. Stone, and R. Lanfear. 2019. The Prevalence and Impact of Model Violations in Phylogenetic Analysis. Genome Biology and Evolution 11:3341–3352.

One Thousand Plant Transcriptomes Initiative. 2019. One thousand plant transcriptomes and the phylogenomics of green plants. Nature 574:679–685.

Philippe, H., H. Brinkmann, D. V. Lavrov, D. T. J. Littlewood, M. Manuel, G. Wörheide, and D. Baurain. 2011. Resolving Difficult Phylogenetic Questions: Why More Sequences Are Not Enough. PLoS Biol 9:e1000602.

Phillips, M. J., and D. Penny. 2003. The root of the mammalian tree inferred from whole mitochondrial genomes. Molecular Phylogenetics and Evolution 28:171–185.

Posada, D. (ed). 2009. Bioinformatics for DNA sequence analysis. Humana Press, New York.

Prum, R. O., J. S. Berv, A. Dornburg, D. J. Field, J. P. Townsend, E. M. Lemmon, and A. R. Lemmon. 2015. A comprehensive phylogeny of birds (Aves) using targeted next-generation DNA sequencing. Nature 526:569–573.

Roycroft, E. J., A. Moussalli, and K. C. Rowe. 2020. Phylogenomics Uncovers Confidence and Conflict in the Rapid Radiation of Australo-Papuan Rodents. Systematic Biology 69:431–444.

Shen, X.-X., D. A. Opulente, J. Kominek, X. Zhou, J. L. Steenwyk, K. V. Buh, M. A. B. Haase, J. H. Wisecaver, M. Wang, D. T. Doering, J. T. Boudouris, R. M. Schneider, Q. K. Langdon, M. Ohkuma, R. Endoh, M. Takashima, R. Manabe, N. Čadež, D. Libkind, C. A. Rosa, J. DeVirgilio, A. B. Hulfachor, M. Groenewald, C. P. Kurtzman, C. T. Hittinger, and A. Rokas. 2018. Tempo and Mode of Genome Evolution in the Budding Yeast Subphylum. Cell 175:1533-1545.e20.

Shen, X.-X., L. Salichos, and A. Rokas. 2016. A Genome-Scale Investigation of How Sequence, Function, and Tree-Based Gene Properties Influence Phylogenetic Inference. Genome Biol Evol 8:2565–2580.

Smith, S. A., N. Walker-Hale, and C. T. Parins-Fukuchi. 2023. Compositional shifts associated with major evolutionary transitions in plants. New Phytologist 239:2404–2415.

Steenwyk, J. L., T. J. Buida, C. Gonçalves, D. C. Goltz, G. Morales, M. E. Mead, A. L. LaBella, C. M. Chavez, J. E. Schmitz, M. Hadjifrangiskou, Y. Li, and A. Rokas. 2022a. BioKIT: a versatile toolkit for processing and analyzing diverse types of sequence data. Genetics 221:iyac079.

Steenwyk, J. L., T. J. Buida, A. L. Labella, Y. Li, X.-X. Shen, and A. Rokas. 2021. PhyKIT: a broadly applicable UNIX shell toolkit for processing and analyzing phylogenomic data. Bioinformatics 37:2325–2331.

Steenwyk, J. L., T. J. Buida, Y. Li, X.-X. Shen, and A. Rokas. 2020. ClipKIT: A multiple sequence alignment trimming software for accurate phylogenomic inference. PLoS Biol 18:e3001007.

Steenwyk, J. L., D. C. Goltz, T. J. Buida, Y. Li, X.-X. Shen, and A. Rokas. 2022b. OrthoSNAP: A tree splitting and pruning algorithm for retrieving single-copy orthologs from gene family trees. PLoS Biol 20:e3001827.

Steenwyk, J. L., Y. Li, X. Zhou, X.-X. Shen, and A. Rokas. 2023. Incongruence in the phylogenomics era. Nature Reviews Genetics, doi: 10.1038/s41576-023-00620-x.

Steenwyk, J. L., G. I. Martínez-Redondo, T. J. Buida, E. Gluck-Thaler, X. Shen, T. Gabaldón, A. Rokas, and R. Fernández. 2024. PhyKIT: A Multitool for Phylogenomics. Current Protocols 4:e70016.

Steenwyk, J. L., D. A. Opulente, J. Kominek, X.-X. Shen, X. Zhou, A. L. Labella, N. P. Bradley, B. F. Eichman, N. Čadež, D. Libkind, J. DeVirgilio, A. B. Hulfachor, C. P. Kurtzman, C. T. Hittinger, and A. Rokas. 2019a. Extensive loss of cell-cycle and DNA repair genes in an ancient lineage of bipolar budding yeasts. PLoS Biol 17:e3000255.

Steenwyk, J. L., and A. Rokas. 2021. orthofisher: a broadly applicable tool for automated gene identification and retrieval. G3 Genes|Genomes|Genetics 11:jkab250.

Steenwyk, J. L., X.-X. Shen, A. L. Lind, G. H. Goldman, and A. Rokas. 2019b. A Robust Phylogenomic Time Tree for Biotechnologically and Medically Important Fungi in the Genera Aspergillus and Penicillium. mBio 10:e00925–19.

Stuart, A. 1955. A TEST FOR HOMOGENEITY OF THE MARGINAL DISTRIBUTIONS IN A TWO-WAY CLASSIFICATION. Biometrika 42:412–416.

Wu, M., J. L. Kostyun, M. W. Hahn, and L. C. Moyle. 2018. Dissecting the basis of novel trait evolution in a radiation with widespread phylogenetic discordance. Molecular Ecology 27:3301–3316.

Yang, Y., M. J. Moore, S. F. Brockington, D. E. Soltis, G. K.-S. Wong, E. J. Carpenter, Y. Zhang, L. Chen, Z. Yan, Y. Xie, R. F. Sage, S. Covshoff, J. M. Hibberd, M. N. Nelson, and S. A. Smith. 2015. Dissecting Molecular Evolution in the Highly Diverse Plant Clade Caryophyllales Using Transcriptome Sequencing. Mol Biol Evol 32:2001–2014.

